# Visualizing cell death in live retina: Using calpain activity detection as a biomarker for retinal degeneration

**DOI:** 10.1101/2021.12.06.471350

**Authors:** Soumaya Belhadj, Nina Hermann, Gustav Christensen, Torsten Strasser, François Paquet-Durand

## Abstract

Calpains are a family of calcium-activated proteases involved in numerous disorders. Notably, previous studies have shown that calpain activity was substantially increased in various models for inherited retinal degeneration (RD). In the present study, we tested the capacity of the *t*-BOC-Leu-Met-CMAC calpain-specific substrate to detect calpain activity in living retina, in organotypic retinal explant cultures derived from wild-type mice, as well as from *rd1* and *Rho*^*P23H/+*^ RD-mutant mice. Test conditions were refined until the calpain substrate readily detected large numbers of cells in the photoreceptor layer of RD retina but not in wild-type retina. At the same time, the calpain substrate was not obviously toxic to photoreceptor cells. Comparison of calpain activity with an immunostaining for activated calpain-2 furthermore suggested that individual calpain isoforms may be active in distinct temporal stages of photoreceptor cell death. Notably, calpain-2 activity may be a relatively short-lived event, occurring only towards the end of the cell death process. Finally, our results support the development of calpain activity detection as a novel *in vivo* biomarker for RD, suitable for combination with non-invasive imaging techniques.

## Introduction

Inherited retinal degeneration (RD) is a group of genetic diseases affecting the retina. They are characterized by progressive photoreceptor degeneration and lead to vision impairment and ultimately blindness. Mutations in at least 280 genes (https://sph.uth.edu/retnet; information retrieved in October 2021) have been associated with RD. The prevalence of monogenic RD is approximately 1 in 3500 individuals (Bertelsen et al. 2014).

RD is largely untreatable and the lack of therapy is due to several reasons: The wide genetic heterogeneity of the disease (Berger, Kloeckener-Gruissem, and Neidhardt 2010), the difficult delivery of therapeutic compounds across the blood retinal barrier (Himawan et al. 2019), and the lack of biomarkers to accurately detect the progression of the disease early on. Typically, RD shows a slow progression with the appearance of symptoms often occurring only in an advanced disease state. Detecting and addressing early phases of pathophysiological cascades is therefore of great importance for therapy development and diagnosis. Moreover, early biomarkers would be of great help to characterize patient populations and to assess treatment efficacy (Hansson 2021).

Most of the mutations causing RD affect genes related to the phototransduction cascade and often cause a dysregulation of cGMP. The toxicity of high levels of cGMP for photoreceptors was already established in the seventies (Farber and Lolley 1974). The two main known targets of cGMP are protein kinase G (PKG) and CNG channels (CNGC). Notably the activity of the latter leads to an increased intracellular calcium concentration (Power, Das, et al. 2020), which may subsequently cause an activation of calcium-activated proteases belonging to the calpain family (Das, Chen, et al. 2021). In this 15-member protease family, calpain-1 and calpain-2, also known as μ-calpain and m-calpain (Goll et al. 2003), are activated by micromoles and millimoles of calcium, respectively (Khorchid and Ikura 2002). While calpain-1 activation mediates synaptic plasticity and neuroprotection, calpain-2 limits the extent of plasticity and may promote neurodegeneration (Baudry 2019; Baudry and Bi 2016).

Incidentally, many RD animal models, with a variety of different disease-causing mutations, display extensive activation of calpains in degenerating photoreceptors. This includes the *rd1, rd2*, and *rd10* mouse models, the *Rho* S334ter, *Rho* P23H, WBN/Kob rat models, as well as photoreceptor degeneration induced by treatment with NaIO_3_ or N-methyl-nitrosourea (MNU) (Arango-Gonzalez et al. 2014; Paquet-Durand et al. 2006; Carido et al. 2014; Tao et al. 2015; Paquet-Durand, Johnson, and Ekström 2007; Power, Rogerson, et al. 2020; Azuma et al. 2004). To assess photoreceptor calpain activity, a well-established assay is available that uses the *t*-BOC-Leu-Met-CMAC (CMAC) calpain substrate on unfixed, *ex vivo* tissue (Ekstrom et al. 2014; Paquet-Durand et al. 2006).

In the present study, we tested the capacity of the CMAC substrate to detect calpain activity in living tissue, in organotypic retinal explant cultures derived from the *rd1, Rho*^*P23H/+*^, and wild-type mice. An evaluation of retinal cytotoxicity using the TUNEL assay indicated that the CMAC substrate was not toxic to photoreceptor cells. Moreover, the CMAC substrate was used to further study the role of calpains in the cell death mechanism leading to retinal degeneration. Overall, the newly established assay procedure is fast and easy to perform and could potentially lend itself to *in vivo* biomarker development.

## Material & Methods

### Animals

C3H HeA *Pde6b*^*rd1/rd1*^ (*rd1*) (Sanyal and Bal 1973), *Rho* ^*P23H/+*^ (Sakami et al. 2011) and wild-type (WT) mice were used. Animals were housed under standard white cyclic lighting, had free access to food and water, and were used irrespective of gender. Animal protocols compliant with §4 of the German law of animal protection were reviewed and approved by the Tübingen University committee on animal protection (Einrichtung für Tierschutz, Tierärztlichen Dienst und Labortierkunde; Registration No. AK02/19M and AK05/19M).

### Organotypic retinal explants

The organotypic retinal explant cultures were prepared as described previously (Belhadj et al. 2020). At post-natal (P) day 5, *rd1* animals were killed and the eyes rapidly enucleated in an aseptic environment. The entire eyes were incubated in R16 serum-free culture medium (Gibco, Carlsbad, CA), with 0.12 % proteinase K (MP Biomedicals, Illkirch-Grafenstaden, France), at 37 °C for 15 min, to allow preparation of retinal cultures with retinal pigment epithelium (RPE) attached. The proteinase K was inactivated with 10 % FCS (Gibco) in R16 medium, and thereafter the eyes were dissected aseptically in a Petri dish containing fresh R16 medium. The anterior segment, lens, vitreous, sclera, and choroid were carefully removed by fine scissors, and the retina was cut perpendicular to its edges, giving a cloverleaf-like shape. Subsequently, the retina was transferred to a culture dish with membrane insert (COSTAR, NY) with the RPE layer facing the membrane. The insert was put into a six-well culture plate and incubated in R16 medium with supplements at 37 °C. The full volume of nutrient medium, 1 ml per dish, was replaced with fresh R16 medium every second day. The procedures for retinal explants cultures derived from *Rho* ^*P23H/+*^ animals were the same as for *rd1* mice, except that the *Rho* mutants were killed at P14 and then cultured for four days until P18.

### TUNEL Assay

The terminal deoxynucleotidyl transferase dUTP nick end labeling (TUNEL) assay (Gavrieli, Sherman, and Ben-Sasson 1992) was performed using the fluorescein kit from Roche Diagnostics (Mannheim, Germany). Sections were incubated in blocking solution (1 % BSA, 10 % normal goat serum, 1 % fish gelatin) for 1 h after 5 min incubation with alcohol acetic acid mixture (62 % EtOH, 11 % Acetic Acid, 27 % H2O). Sections were then stained using the kit as per manufacturer’s instructions.

### Immunohistochemistry

The fixed cryosections were dried at 37°C for 30 min and washed in PBS for 10 min. Then, blocking solution (10 % NGS and 1 % BSA in 0.3 % PBST) was applied to the tissue and incubated for 1 h at room temperature (RT). The anti-calpain-2 primary antibody (Abcam, Cambridge, UK) was diluted 1:300 in blocking solution and incubated with the sections overnight at 4°C. The sections were then rinsed with PBS three times for 10 min each. The secondary antibody (goat anti-rabbit Alexa Fluor 488 or 568, Molecular Probes, Eugene OR) was diluted 1:500 in PBS and incubated with the sections for 1 h at RT, in the dark. The sections were rinsed three times with PBS for 10 min each, mounted with and stored in 4 °C for at least 30 min before imaging.

### Detection of calpain activity in retinal explants

The cultures derived from the *rd1* mouse model were treated for variable incubation periods (1 h, 3 h, 6 h, and 24 h) with the substrate tertiary-butyloxycarbonyl-leucine-methionine-7-amino-4-methylcoumarin (abbreviated to *t*-BOC-Leu-Met-CMAC, or in short: CMAC) (Invitrogen, Carlsbad, CA), at a concentration of 50 μM in the culture medium. CMAC was diluted in the culture medium at P11 for a 24 h incubation or at P12 for the other conditions (1 h, 3 h, and 6 h). At the end of the incubation time, the cultures were stopped by 45 min fixation in 4 % PFA. This was followed by graded sucrose cryoprotection, embedding in Tissue-Tek O.C.T. compound (Sakura Finetek Europe, Alphen aan den Rijn, Netherlands)-filled boxes, and collecting of 12-μm-thick retinal cross-sections on a Thermo Scientific NX50 cryotome (Fig. 1).

**Figure 1:**
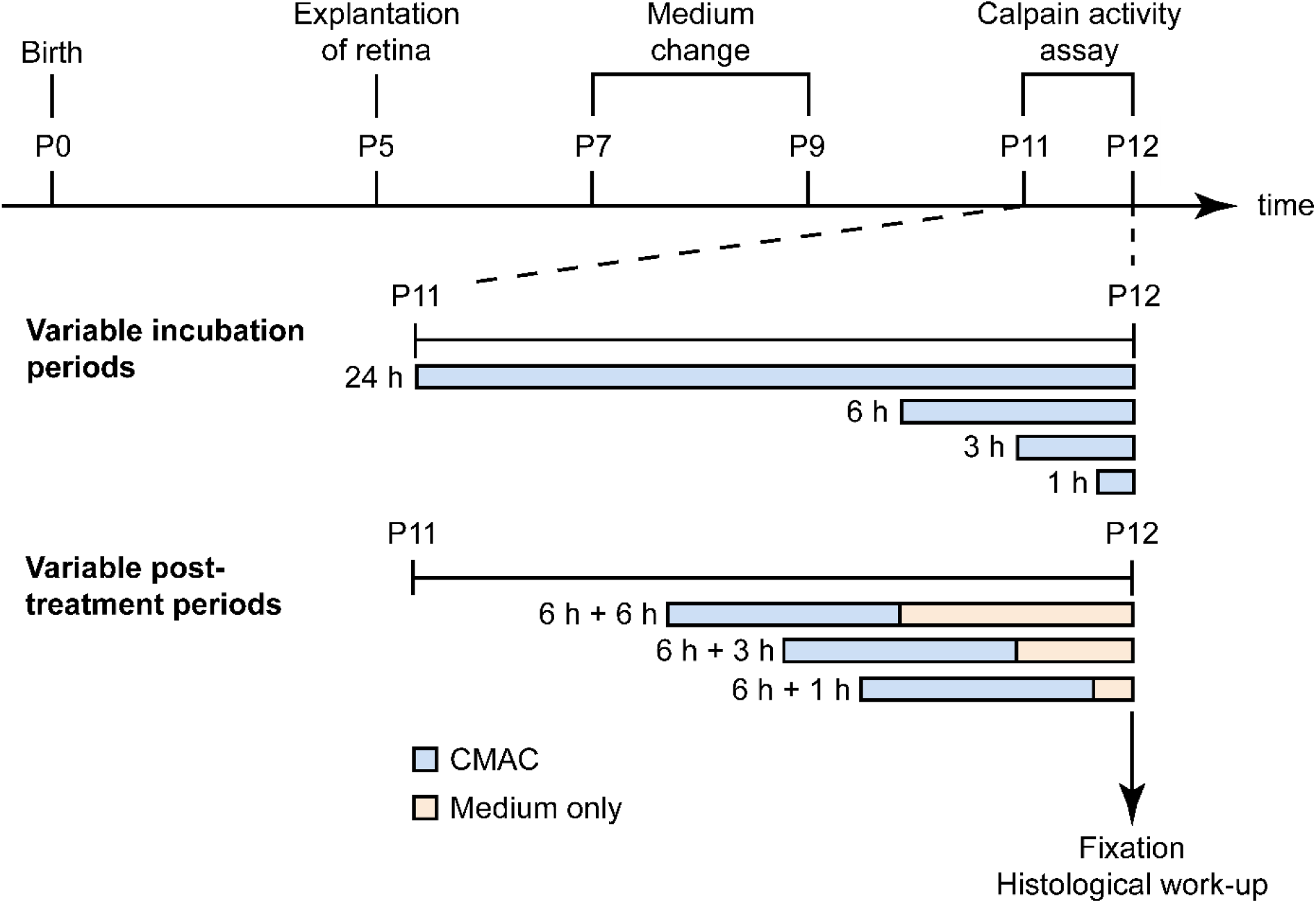
Overview of experimental paradigms for calpain activity detection in retinal explants. The organotypic retinal explant cultures were derived from either wild-type or RD mutant animals at post-natal day (P) 5. The culture medium was changed every second day and the retinal explants were cultivated until P12. In the first set of experiments the CMAC substrate was diluted in the culture medium between P11 and P12 and incubated for variable periods, ranging from 1 h to 24 h. In the second experimental paradigm the retinal explants were incubated with CMAC substrate for 6 h, then the medium was replaced by fresh culture medium without CMAC substrate, and the incubation was continued for another 1 h, 3 h, or 6 h. At P12 the cultures were ended by fixation in 4 % PFA.

In a second set of experiments the period of time that a given cell would remain positive for calpain activity was investigated. To this end, retinal cultures were incubated at P12 for 6 h with the CMAC substrate (Invitrogen, Carlsbad, CA), at a concentration of 50 μM in the culture medium. Thereafter, the medium containing the CMAC substrate was replaced by fresh culture medium for different incubation times: 1 h, 3 h, 6 h. Afterwards, the cultures were stopped by 45 min fixation in 4 % PFA and further processing as above. The cultures derived from the *Rho*^*P23H/+*^ mouse model were treated with the CMAC substrate at P18 for 6 h at a concentration of 50 μM in the culture medium. Afterwards, the cultures were stopped by 45 min fixation in 4 % PFA and further processing as above (Fig. 1).

### Microscopy, cell counting, and statistical analysis

Light and fluorescence microscopy were performed at room temperature on an Axio Imager Z.1 ApoTome Microscope, equipped with a Zeiss Axiocam MRm digital camera (Zeiss, Oberkochen, Germany). Images were captured using the Zeiss Zen software; representative pictures were taken from central areas of the retina using a 20x / 0.8 Zeiss Plan-APOCHROMAT objective. For quantifications, pictures were captured on three entire sagittal sections from at least three different animals. The average area occupied by a photoreceptor cell (*i.e*., cell size) was determined by counting 4’
s,6-diamidino-2-phenylindole (DAPI) stained nuclei in nine different areas of the retina. The total number of photoreceptor cells was estimated by dividing the outer nuclear layer (ONL) area by this average cell size. The number of positively labelled cells in the ONL was counted manually. Errors in graphs and text are given as standard deviation (STD).

The data on calpain activity positive cells in the ONL after variable post-treatment periods was used to fit a one phase exponential decay model (N(t) = N_0_e^-λt^) using GraphPad Prism (GraphPad software, San Diego, CA, USA). This resulted in a baseline value N_0_ of 7.097 and a decay rate λ of 1.630 (R^2^ = 0.6087). A half-life time for calpain positivity of 0,4252 hours (95 % CI: 0 – 2.2 hours) was calculated as ln(2)/λ (see model in red; Fig. 6E, F).

## Results

### Calpain activity detection in live retina

Calpain activity has previously been assessed *ex vivo* on unfixed retinal tissue sections using CMAC (Fig. 2) (Arango-Gonzalez et al. 2014; Ekstrom et al. 2014; Paquet-Durand et al. 2006). Here, we tested whether CMAC could also be used to detect calpain activity in living organotypic retinal explant cultures. The CMAC substrate is processed by endogenous glutathione S-transferase which conjugates glutathione to the coumarin segment of the molecule (Habig, Pabst, and Jakoby 1974; Rosser, Powers, and Gores 1993). Then, the cleavage by calpain results in red-shifting and enhancement of the fluorescence emission peak of the MAC-SG product. CMAC is hydrophobic and thus potentially suitable for calpain activity detection *in vivo* as it is expected to cross the relevant biological membranes like the outer blood retinal barrier (Pitkanen et al. 2005) and cell membrane (Rosser, Powers, and Gores 1993). After glutathione conjugation, the resulting MAC-SG product is considered to be cell impermeable (Rosser, Powers, and Gores 1993).

**Figure 2:**
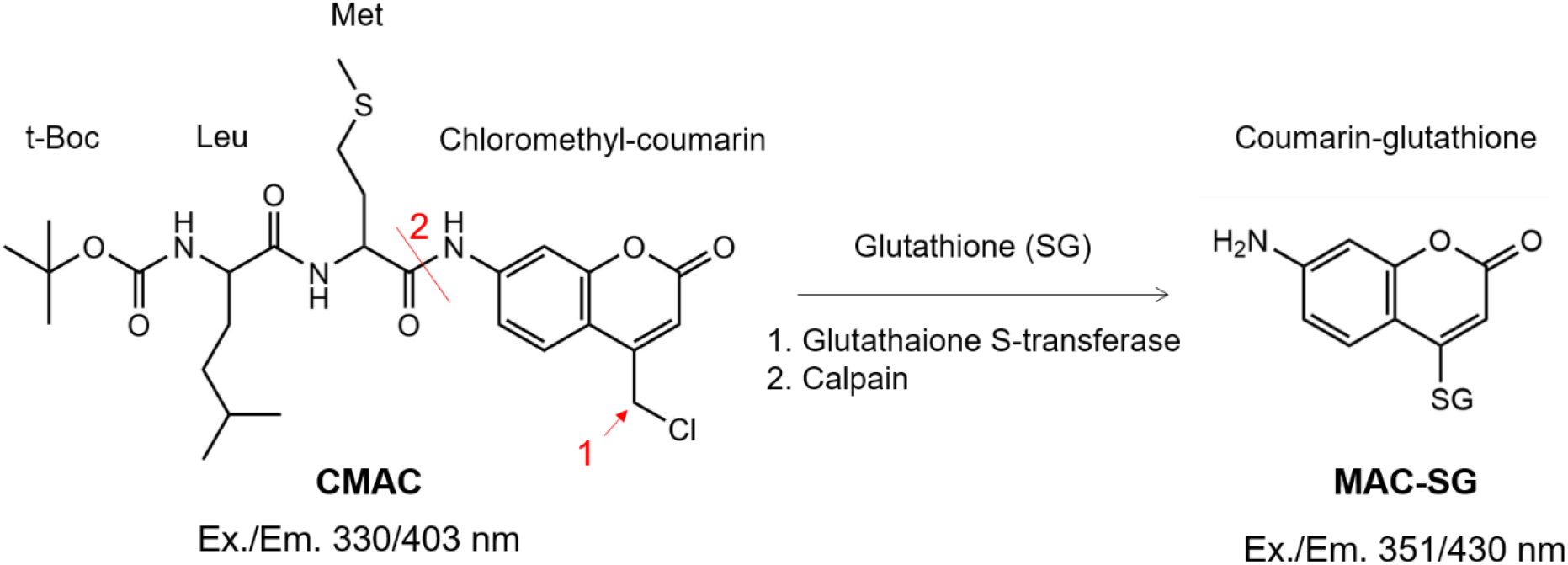
Molecular structure of the calpain substrate CMAC and product MAC-SG. The calpain activity assay with the CMAC substrate involves an initial glutathione conjugation to the coumarin segment at position indicated by the arrow 1. Afterwards, calpain cleaves the peptide bond at position 2, unquenching the coumarin-glutathione product and red-shifting the excitation and emission peaks.

To test whether the CMAC substrate could be used for the detection of calpain activity in living tissue, we used organotypic retinal explant cultures and exposed them to CMAC for 1 h, 3 h, 6 h, and 24 h (Fig. 3A-D). Retinal explants were derived from either wild-type mice or from two distinct RD models, the *rd1* and *Rho*^*P23H/+*^ mouse.

**Figure 3:**
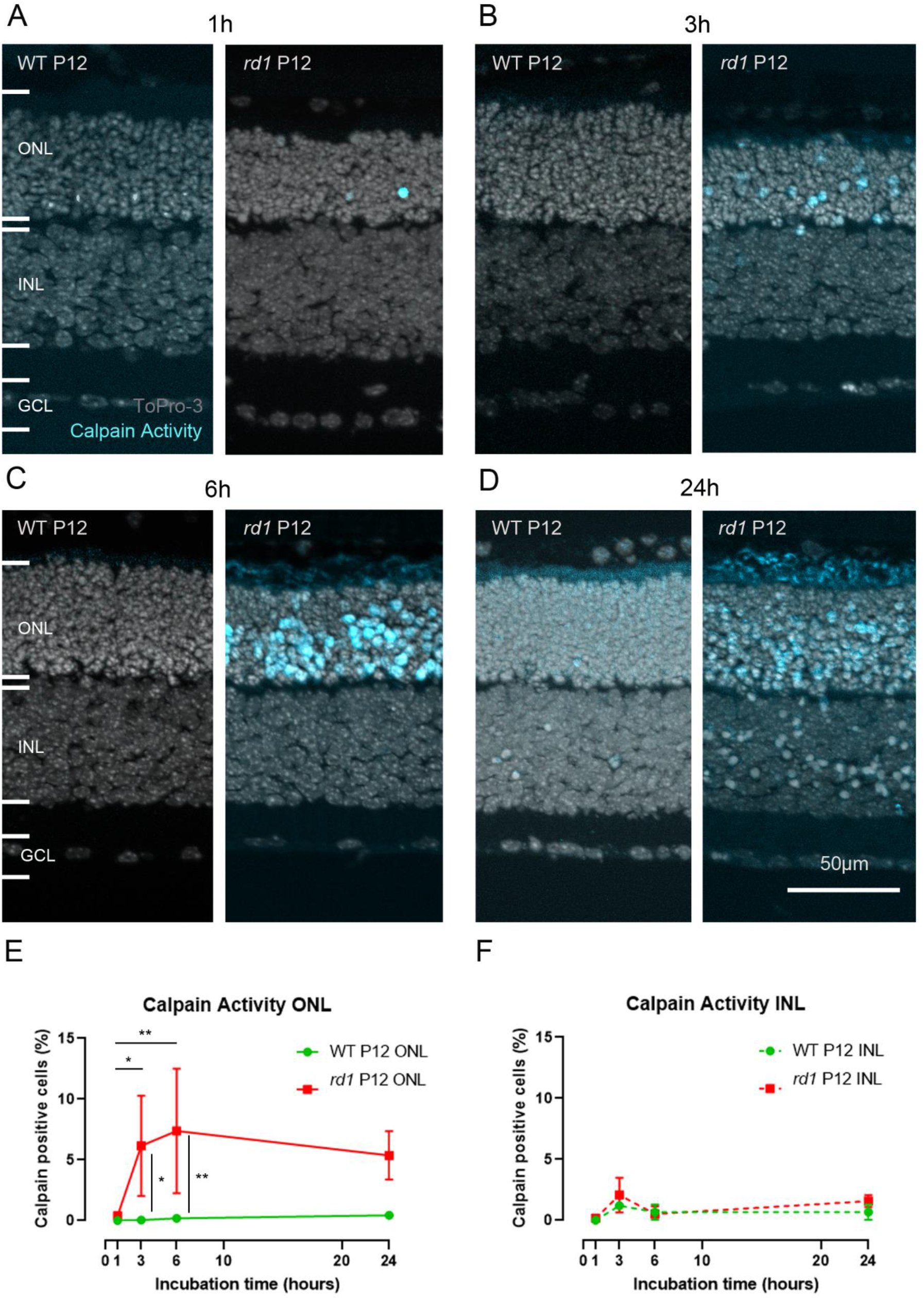
Live tissue detection of calpain activity in the *rd1* mouse model. Organotypic retinal explant cultures were incubated with 50 μM CMAC for 1 h (**A**); 3 h (**B**); 6 h (**C**); 24 h (**D**) before the culture was ended by fixation. To-Pro-3 was used as nuclear counterstain. The number of calpain positive cells detected in the *rd1* ONL increased from 1 h to 6 h to finally reach a plateau (**E**), while in the INL no specific trend was observed (**F**). Images representative for results obtained from at least 3 independent retinal explant cultures; error bars indicate STD; statistical analysis: two-way ANOVA; * = *p* < 0.05; ** = *p* < 0.01; GCL = ganglion cell layer.

In *rd1* mouse retinal explants, an increasing number of calpain active photoreceptors (*i.e*., cells in the outer nuclear layer, ONL) was detected when the CMAC incubation time was increased from 1 h to 6 h (Fig. 3A, B, C, E), with respectively 0.4 % (± 0.3, n = 4) and 7.4 % (± 5.1, n = 4) detected positive cells. After 6 h of incubation, the calpain active cell detection rate appeared to no longer increase. Indeed, after 24 h incubation, the amount of detected positive cells was 5.3 % (± 1.9, n = 4), which was not significantly different from the previous incubation time (6 h) (Fig. 3D, E). The numbers of detected calpain positive cells in the ONL between the 6 h and 24 h time points corresponded to what was observed in *ex vivo* retinal sections from the *rd1* animal model at this age (Ekstrom et al. 2014; Paquet-Durand et al. 2006; Power, Rogerson, et al. 2020). In the inner nuclear layer (INL), the number of detected calpain active cells was significantly lower than in the ONL for both the *rd1* and wild-type conditions (Fig. 3A, B, C, D, F). In wild-type explants, the number of calpain positive cells remained low throughout the incubation period.

To address the question whether calpain activity could also be detected in live retina suffering from a dominant mutation in the rhodopsin gene, we furthermore investigated explant cultures derived from heterozygous *Rho*^*P23H/+*^ mice. According to previous studies on this model, rod photoreceptor degeneration starts around P14, peaks at P18-P20 and is almost completed by P31 (Comitato et al. 2020; Nakamura et al. 2017). At P18, calpain activity could also be detected in the *Rho*^*P23H/+*^ mouse model (Fig. S2B, C), although the extent of calpain activity was comparatively low with 1.09 % (± 0.7, n = 3) positive cells in the ONL. At the same time the number of dying, TUNEL positive cells was 3.8 % (± 0.7, n = 3) (Fig. S2A, C).

### The calpain activity assay exhibits low toxicity in live retina

Since the CMAC substrate had so far not been tested in live retinal cultures, the potential toxicity of the compound was investigated. Organotypic retinal explant cultures derived from *rd1* and wild-type mice were treated either with the CMAC substrate or without (2.5 % DMSO) at different incubation times: 1 h, 3 h, 6 h, and 24 h. The TUNEL assay (Gavrieli, Sherman, and Ben-Sasson 1992) was then used for the detection of cell death on fixed tissue sections from these cultures.

Qualitatively, it was observed that the typical histology of the retina was consistently preserved after all treatments (Fig. 3A-D & 4A-D). Additionally, the number of TUNEL positive cells detected in the ONL was not significantly different between the different incubation periods for *rd1* and wild-type animals (Fig. 4E), with approximately 2.7 – 2.8 % detected positive cells in the *rd1* model across incubation times and from 1.0 to 1.6 % in the wild-type. Also, the number of TUNEL positive cells detected in the ONL is in accordance with what is usually observed in these models at the same age (Arango-Gonzalez et al. 2014; Ekstrom et al. 2014; Power, Rogerson, et al. 2020).

**Figure 4:**
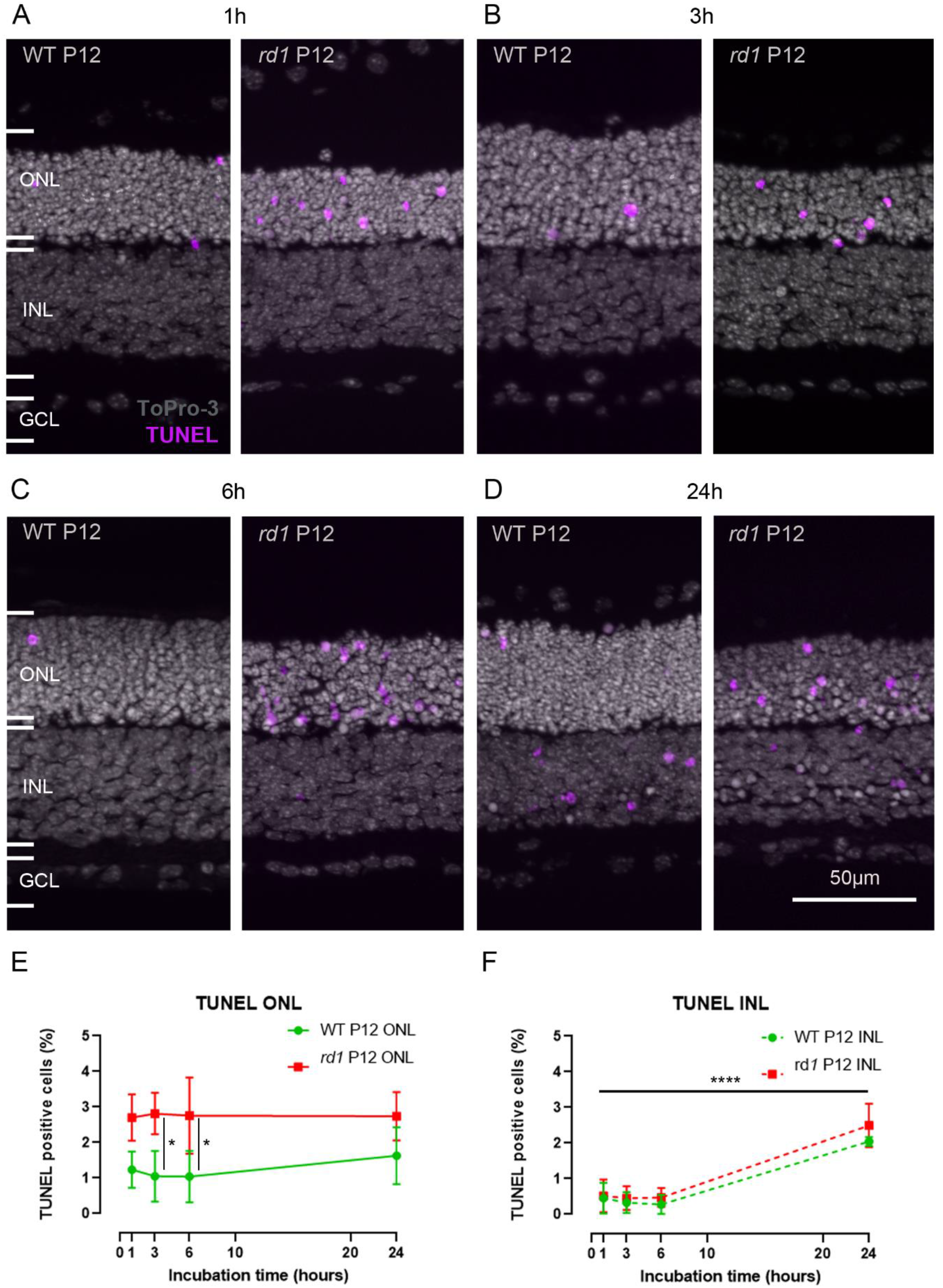
Cell death in live tissue treated with the CMAC substrate. Organotypic retinal explant cultures derived from the *rd1* mouse model were incubated with 50 μM CMAC for 1 h (A); 3 h (B); 6 h (C); 24 h (D). A TUNEL assay for cell death detection was carried out on fixed tissue sections. Percentage of TUNEL positive cells in the ONL (E) and the INL (F). To-Pro-3 was used as nuclear counterstaining.Images are representative for results obtained from at least 3 independent retinal explant cultures; error bars indicate STD; statistical analysis: two-way ANOVA. ONL = outer nuclear layer, INL = inner nuclear layer, GCL = ganglion cell layer.

In the INL, the number of TUNEL positive cells remained insignificant in cultures treated with the CMAC substrate from 1 h to 6 h, with approximately 0.5 % positive cells detected in the *rd1* model and approximately 0.3 % positive cells detected in the wild-type. However, an increase in cell death was observed when cultures derived from both *rd1* and wild-type mice were treated with the CMAC substrate for 24 h with 2.5 % (± 0.6, n = 3) positive cells detected in the *rd1* model and 2.0 % (± 0.1, n = 3) positive cells detected in the wild-type (Fig. 4F). This increase in the TUNEL counts was not observed after the same incubation time with the vehicle (DMSO, 2.5 %) (Fig. S1), suggesting that the observed effect could be due to prolonged exposure to the CMAC substrate.

### Calpain activity and TUNEL assay partly colocalize

To further relate calpain activity with cell death, we studied the colocalization of TUNEL positive, dying cells with the calpain activity assay. In the ONL around 3.8 % (± 1.2, n = 3) of cells were found to be TUNEL positive (Fig. 5), which is in accordance with previous studies (Sahaboglu et al. 2016). Approximately 6.2 % (± 0.6, n = 3) of photoreceptors stained positive for the calpain substrate CMAC, which was used as a proxy for overall calpain activity (Fig. 5A, B, C). Quantification of colocalization between TUNEL and calpain activity showed that 1.3 % (± 0.5, n = 3) of all cells were double-positive for both assays (Fig. 5C, D). Hence, only about 20% of the cells exhibiting calpain activity also showed TUNEL positivity. Since the TUNEL assay is thought to label cells at the very end of the cell death process (Sahaboglu et al. 2013), this implies that around 80 % of calpain activity positive cells had still not reached the final stages of cell death.

**Figure 5:**
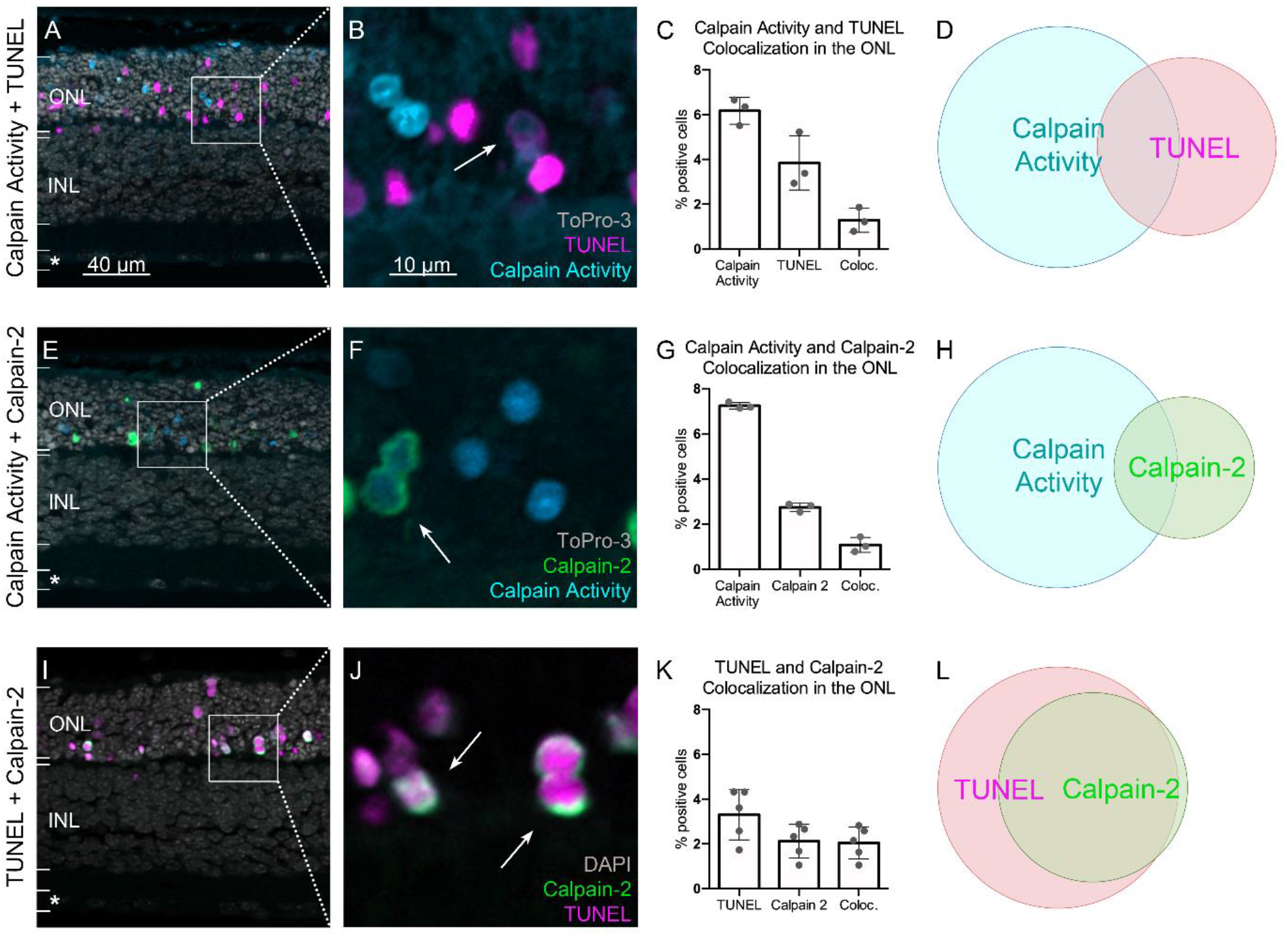
Colocalization of the TUNEL assay, calpain activity assay, and calpain-2 immunostaining. (**A-D**) Comparison between calpain and calpain-2 activation. (**E-H**) Comparison between calpain activity and the TUNEL assay. (**I-L**) Comparison between calpain-2 activation and the TUNEL assay. The arrows highlight cells which show colocalization. **D, H**, and **L** show Venn diagrams, which visualize the relative proportion of cells which are positive for the respective staining as well as the overlap between different stainings (colocalization). Images representative for results obtained from 3-5 independent retinal explant cultures derived from the *rd1* mouse model. Error bars indicate STD. ONL = outer nuclear layer, INL = inner nuclear layer, asterisk = ganglion cell layer.

The calpain family comprises 15 isoforms and the CMAC substrate is not specific to a particular isoform. Since calpain-2 had previously been reported as the isoform driving neurodegeneration (Baudry 2019; Baudry and Bi 2016), we performed an immunostaining for activated calpain-2 to assess the involvement of this isoform in the neurodegenerative process. While in this situation 7.2 % of the cells in the ONL (± 0.1, n = 3) were detected as calpain positive, only 2.7 % (± 0.2, n = 3) of these cells stained positive for activated calpain-2 (Fig. 5E, F, G). Regarding the colocalization between calpain activity and activated calpain-2, 1 % (± 0.3, n = 3) of all cells were double positive (Fig. 5G, H). The staining of activated calpain-2 presented as a ring along the periphery of photoreceptor cell bodies, which corroborates the finding that calpains are activated at the cell membrane upon the availability of Ca^2+^ and phospholipids (Suzuki et al. 2004) (Fig. 5F, J).

A further colocalization experiment was carried out between TUNEL and activated calpain-2. While 3.3 % (± 1.1, n = 5) of photoreceptors were positive for TUNEL, 2.1 % (± 0.8, n = 5) of them were positive for activated calpain-2 (Fig. 5I, J, K). Around 2 % (± 0.7, n = 5) of all photoreceptors were double-positive for TUNEL and activated calpain-2, which accounts for almost all cells positive for activated calpain-2 (Fig. 5K, L). This strong overlap between TUNEL and calpain-2 activation indicated that calpain-2 activation occurred only in the final stages of the cell death process.

### Estimating the duration of calpain activity in individual dying photoreceptors

To assess for how long the activity identified with the CMAC substrate (MAC-SG) was detectable in single ONL cells, we first treated *rd1* retinal explants with CMAC and then reverted to unlabeled culture medium without CMAC for post-treatment incubation periods ranging from 0 to 6 h (Fig. 6A-D). Here, the retinal cultures were treated with the CMAC substrate for 6 h as this was the incubation time for which we observed the strongest signal (*cf*. Fig. 3).

**Figure 6:**
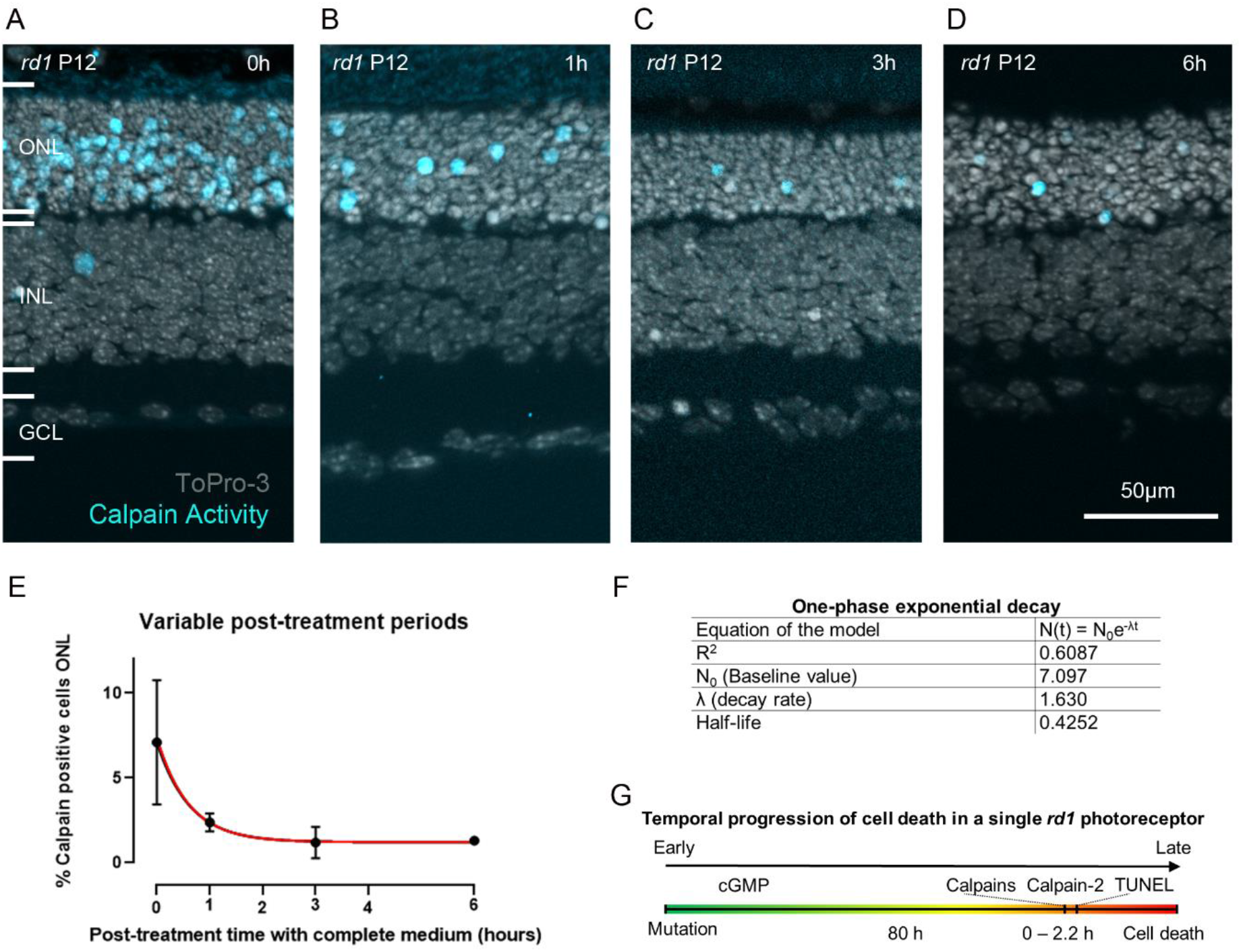
Estimation of calpain activity duration in photoreceptors during the cell death process. (**A-D**) Organotypic retinal explant cultures derived from the *rd1* mouse model were treated with the CMAC substrate for 6 h. The medium containing CMAC was then replaced by plain medium for varying durations: 0 h (**A**), 1 h (**B**), 3 h (**C**), 6 h (**D**). (**E**) Quantification of calpain positive cells in the ONL. The number of detected calpain positive cells over time followed an exponential decay. Red line shows model fit. (**F**) Table showing model equation and parameters. (**G**) Putative timeline for disease progression in an individual photoreceptor cell. Error bars indicate STD. INL = inner nuclear layer, GCL = ganglion cell layer. Images representative for results obtained from 4 independent retinal explant cultures per condition.

The amount of detected calpain positive cells decreased sharply with the post-treatment incubation period until it reached a lower threshold at approximately 3 h post-treatment. Indeed, at 0 h post-treatment incubation period, 7.4 % (± 5.1, n = 4) calpain positive cells were detected, while 2.4 % (± 0.5, n = 4), 1.2 % (± 0.9, n = 4) and 1.3 % (± 0.2, n = 3) calpain positive cells were detected after 1 h, 3 h, and 6 h post-treatment incubation periods, respectively (Fig. 6A-F). Based on the logic forwarded in a study by Clarke and colleagues (Clarke et al. 2000), we fitted a one phase exponential decay model to the calpain activity data. The model estimates for the half-life time of a calpain positive cell were ranging between 0 – 2.2 hours, with a best fit prediction of around 0.4 hours. In other words, in a given dying photoreceptor cell it will likely take a little less than 30 minutes for the calpain activity signal to dissipate.

## Discussion

The detection and quantification of cellular calpain activity is highly desirable in the context of many diseases, including neurodegenerative diseases of the retina. In this study, we used the CMAC substrate to monitor calpain activity in living retina and to relate it to cell death and the activity of the calpain-2 isoform. Since the retina is accessible to non-invasive *in vivo* imaging, using, for instance, scanning laser ophthalmoscopy, the direct detection of calpain activity appears feasible and could significantly improve our understanding of physiological and pathological processes. Moreover, calpain activity detection could serve as a surrogate biomarker for the study and diagnosis of neurodegenerative diseases.

### Detection of calpain activity

Numerous assay kits using fluorogenic or luminogenic specific calpain substrates are commercially available to detect calpain activity in cell or tissue lysates. However, these assays do not offer cellular resolution of calpain activity (Tian et al. 2020). Calpain activity can be detected indirectly by using antibodies recognizing substrate proteins after calpain specific cleavage. For example, antibodies recognizing the 150 kDa cleaved fragment of α-fodrin have been used to detect calpain-specific fragmentation using the Western blot technique or immunofluorescence (Lee et al. 2018; Saatman et al. 2003). Upon Ca^2+^ dependent activation calpains undergo an autolytic cleavage that reduces their apparent molecular weight by about 2 kDas (Suzuki et al. 2004). This opens another way to assess calpain activity on histological tissue sections via the use of antibodies recognizing only the activated calpain-1 or calpain-2 protease (Power, Rogerson, et al. 2020). Moreover, *in vivo* imaging of calpain activity is enabled in a mouse model which ubiquitously expresses a FRET reporter consisting of eCFP and eYFP, and separated by a linker cleavable by calpains (Bartoli et al. 2006; Stockholm et al. 2005). As opposed to these approaches, the detection method forwarded in this study opens the possibility to perform *in vivo* imaging of calpain activity without the use of transgenic animals or antibody detection systems.

The apparent sensitivity of the live tissue assay, as determined by the positive cell detection rate, is similar to that of the established assay using the same substrate on unfixed dead tissue sections (Ekstrom et al. 2014; Paquet-Durand et al. 2006). Moreover, calpain activity could be detected in organotypic retinal explants derived from two genetically distinct mouse models for retinal degeneration, suggesting a general applicability of the assay in the genetically very heterogeneous RD-type diseases. The mutation underlying the *rd1* mouse phenotype affects the *Pde6b* gene. In humans, PDE6B mutations are responsible for retinitis pigmentosa (RP) in about 4-5 % of patients, following an autosomal-recessive way of inheritance (Berger, Kloeckener-Gruissem, and Neidhardt 2010). Mutations in the rhodopsin gene are the most common form of autosomal-dominant retinitis pigmentosa (adRP) and the variant RHO^P23H^ is the most common cause of adRP in the United States (Sohocki et al. 2001). It is interesting to note that in the *rd1* model, at P11, *i.e*. at the onset of degeneration, calpain active cells numbered about twice as many as TUNEL positive cells. In the *Rho*^*P23H/+*^ model, at P18, a comparable time-point in the progression of retinal degeneration, the calpain positive cells were outnumbered by TUNEL positive cells in a ratio of about 1 to 4. This discrepancy between the recessive *rd1*- and dominant *Rho*^*P23H/+*^ models could point at difference in the execution of cell death, perhaps associated with different kinetics of the various processes that govern the degeneration.

### The calpain activity assay exhibits low toxicity in live retinal explant cultures

The fact that the histology of the tissue remained undisturbed after the different incubation times with the CMAC substrate, as well as the TUNEL count in ONL and INL for incubation times up to 6 h, suggests that the CMAC substrate is not toxic to photoreceptors and to the retina in general. This is in line with cell culture-based studies using the CMAC substrate, which also do not report toxicity of the probe (Marzia et al. 2006; Su et al. 2010; Tauskela et al. 2000). The increase in TUNEL counts in the INL after the 24 h incubation time with CMAC in both *rd1* and wild-type mice suggests that the substrate could be toxic to INL cells after prolonged exposure. Future studies using specific markers for different INL cell types may reveal whether certain cell populations might be sensitive to CMAC.

### Calpain activity: a short event in the photoreceptor death process?

The fluorescent signal from the CMAC calpain substrate was extinguished in a time-course that appeared to follow an exponential decay. Indeed, previous studies suggest that cell death follows exponential kinetics (Clarke et al. 2000). The half-life for calpain positivity, estimated to be between 0 and 2.2 hours (95 % CI), may seem relatively short given that the total duration of photoreceptor cell death in *rd1* retina was estimated to last around 80 hours (Sahaboglu et al. 2013). Still, since the MAC-SG product is considered to be cell impermeable (Rosser, Powers, and Gores 1993), it is unlikely that the extinction of the signal was caused by diffusion out of the cell. Instead, the duration of calpain activity is likely a rather short event during the photoreceptor cell death process.

We observed that even less cells were positive for activated calpain-2 than for overall calpain activity. Since the CMAC labelling captures the activity of all calpain isoforms present in the retina, including but not exclusive to calpain-2, this discrepancy is to be expected. Furthermore, the calpain activity assay was performed on live retinal explants over a duration of 6 hours, while the calpain-2 staining captures a snapshot of the cells in which calpain-2 was activated precisely at the time-point of fixation. Another reason might be that calpain-2 is inactivated very quickly via autolysis (Hata et al. 2001; Moldoveanu et al. 2003), which might lead to only small amounts of CMAC being cleaved by calpain-2. Further co-staining experiments, including other calpain isoforms and other signaling molecules involved in cGMP-dependent cell death, could help to unravel the time course of photoreceptor degeneration. Due to a lack of available antibodies and the lethality of certain calpain knock-out models (Dutt et al. 2006; Takano et al. 2011), this may require the development of appropriate methodology.

### Calpain-2 is activated in the final stages of photoreceptor cell death

Several recent studies have shown that individual calpain species may serve distinct roles *in vivo* (Baudry 2019; Franco, Perrin, and Huttenlocher 2004; Santos et al. 2012). Furthermore, calpain isoforms show differences regarding the time window of proteolysis, the pattern of autolysis and the concentration of Ca^2+^ required for activation (Shinkai-Ouchi et al. 2020; Yoshimura et al. 1983). Notably, calpain-2 requires 2-3 magnitudes more Ca^2+^ compared to calpain-1 to be activated. The calpain activity as identified by CMAC labeling and calpain-2 immunostaining have previously been associated with cell death in photoreceptor degeneration (Paquet-Durand et al. 2006; Paquet-Durand et al. 2010; Power, Rogerson, et al. 2020). However, the colocalization analysis revealed significant differences between the TUNEL assay, CMAC labeling, and calpain-2. This suggested that the different assays capture dying cells at distinct time-points during the progression of photoreceptor cell death. The TUNEL assay is thought to label cells at the very end of the process of degeneration (Sahaboglu et al. 2013), while calpain activity is believed to occur at a slightly earlier stage (Power, Das, et al. 2020; Power, Rogerson, et al. 2020). The low percentage of co-staining between calpain activity and TUNEL observed here suggested that calpain activity occurred distinctly earlier in the timeline of cell death than TUNEL positivity. Interestingly, the overlap between calpain-2 and calpain activity was also relatively low, suggesting that activation of calpain-2 occurred at a later time interval as compared to overall calpain activity. The very high, but not complete, co-localization between TUNEL and calpain-2 further suggests that calpain-2 activation occurred at a late, nearly final stage of cell death. Together with the fact that calpain-2 activation was restricted to the perinuclear areas of photoreceptors, this raises the question as to how calpain-2 might be activated. Here, a recent study pointed to a potential role of voltage-gated Ca^2+^ channels (VGCC), which are expressed in photoreceptor cell bodies, and which might mediate the influx of Ca^2+^ ions required for calpain-2 activation (Das, Popp, et al. 2021). Combined with the higher availability of Ca^2+^ in the cytosol towards an advanced stage of cell death, due to a degradation of cell organelles and overall dysregulation of Ca^2+^ homeostasis, this might explain the delayed occurrence of calpain-2 activity.

## Conclusion and outlook (CMAC Substrate as a biomarker)

The results presented in this study support a further development of the CMAC substrate as a biomarker for cell death detection. While pharmacodynamics, pharmacokinetics, and toxicity of this calpain substrate need to be further investigated, in the retina, this method could potentially enable a direct, non-invasive detection of calpain activity *in vivo*, with single-cell resolution, using techniques such adaptive optics scanning laser ophthalmoscopy (AO-SLO) (Akyol et al. 2021). Here, an infrared two-photon excitation system could be used to avoid UV-light stimulation. Alternatively, a calpain specific peptidic substrate could be combined with a near infrared fluorophore to enable single-photon stimulation. Taken together, our results suggest that the role of calpain in the process of photoreceptor cell death may be more complex than previously assumed, with individual isoforms being active in slightly distinct temporal stages of cell death. Other than biomarker applications, understanding the temporal progression of calpain activities during photoreceptor cell death may also have direct implications on the development of therapeutic approaches for RD-type diseases.

## Authors contribution statement

SB planned the experiments, collected and analyzed the data, drafted the manuscript and prepared the figures. SB also co-supervised NH during her lab rotation. NH contributed with immunostainings, she performed the experiments, collected and analyzed data, prepared a figure and contributed in writing. GC prepared 2 figures and contributed in writing. TS contributed with data analysis and help and advice with modelling. FPD was involved in supervision, experimental design and data interpretation. All the authors edited the manuscript and provided critical feedback.

## Acknowledgments

We thank Norman Rieger for excellent technical assistance and Mathias Seeliger, Timm Schubert, Per Ekström, and John Groten for helpful discussions. We thank Blanca Arango-Gonzalez for kindly providing the *Rho*^*P23H/+*^ mice and Lucas Kook for his help and insights into modelling.

