## Supplemental Table S1 and Supplemental Figure S1 for "Visualizing cell death in live retina: Using calpain activity detection as a biomarker for retinal degeneration"

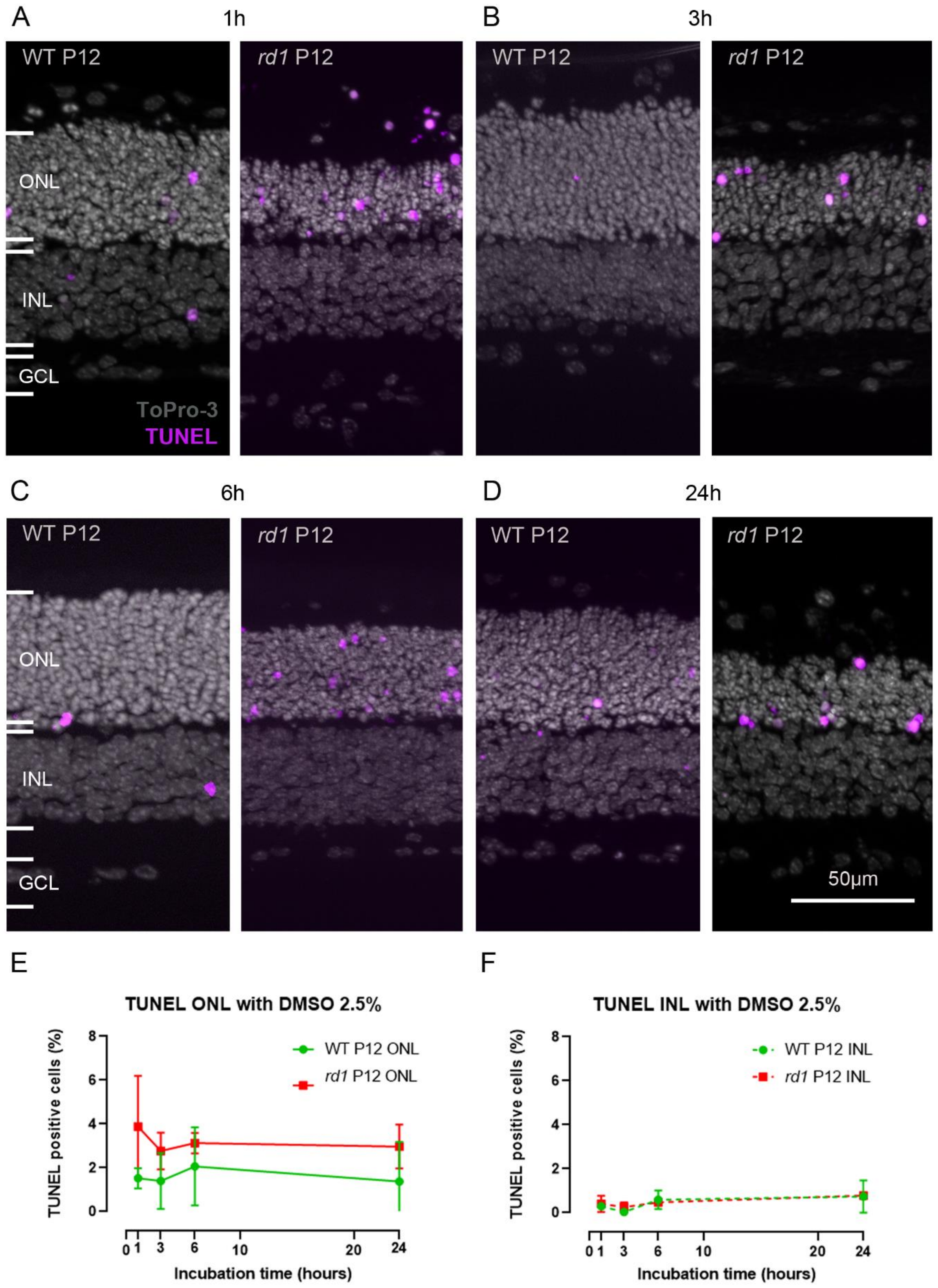


**Figure S1: Cell death in live tissue treated with 2.5 % DMSO (vehicle).** The TUNEL assay (magenta) for cell death detection was carried out on fixed retinal sections, To-Pro-3 (grey) was used as nuclear counterstaining. Organotypic retinal explant cultures derived from the *rd1* mouse model were incubated with: (**A**) DMSO (2.5 %) for 1 h, (**B**) 3h, (**C**) 6 h, (**D**) and 24 h. (**E**, **F**) Percentages of TUNEL positive cells in the outer nuclear layer (ONL) and inner nuclear layer (INL). Images are representative for results obtained from at least 3 independent retinal explant cultures; error bars indicate STD; statistical analysis: two-way ANOVA, GCL = ganglion cell layer.

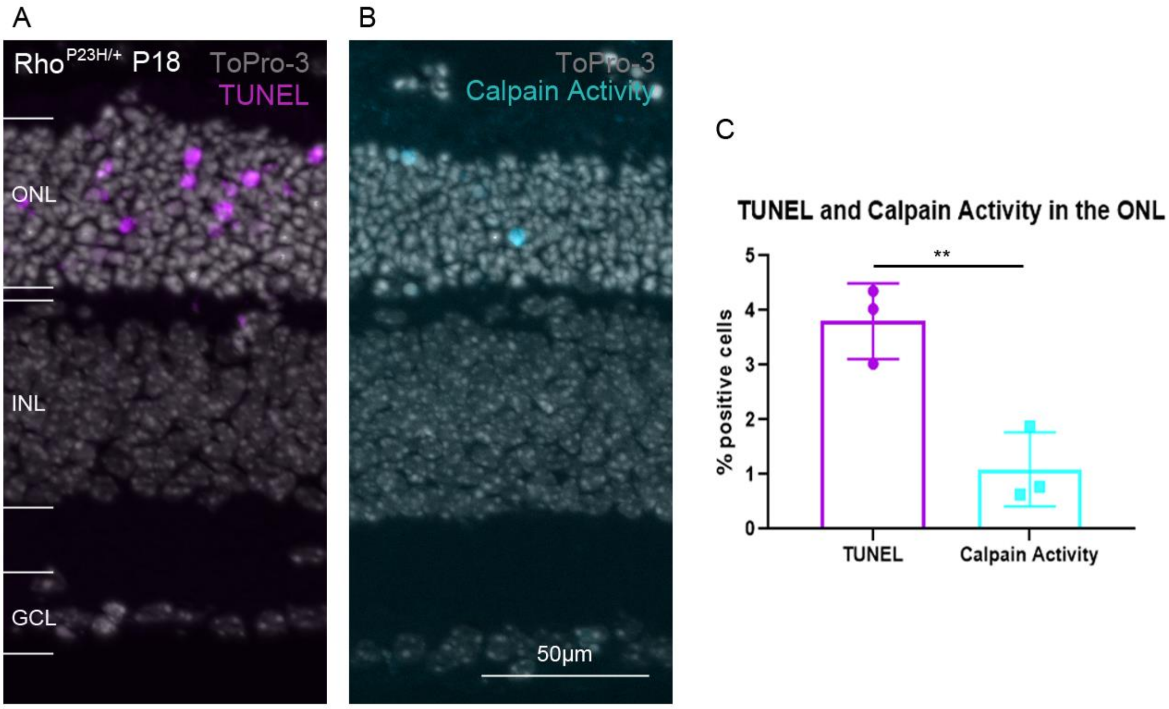


**Figure S2: Live tissue detection of calpain activity and cell death in the *Rho*$^{P23H/+}$ mouse model.** (**A**) The TUNEL assay (magenta) for cell death detection was carried out on fixed tissue sections from organotypic retinal explant cultures derived from the *Rho*$^{P23H/+}$ mouse model. (**B**) Live, organotypic retinal explant cultures were incubated with 50 µM CMAC for 6 h at P18. A calpain specific fluorescent signal (cyan) was observed in individual photoreceptor cells in the outer nuclear layer (ONL). To-Pro-3 (grey) was used as nuclear counterstaining. Images representative for results obtained from at least 3 independent retinal explant cultures; error bars indicate STD; statistical analysis: unpaired t-test. INL = inner nuclear layer, GCL = ganglion cell layer.
